# The dynamics of HD-ZIP III - ZPR protein interactions play essential roles in embryogenesis, meristem function and organ development

**DOI:** 10.1101/2021.11.24.469949

**Authors:** Anna Vitlin Gruber, Melissa Kosty, Yasaman Jami-Alahmadi, James A. Wohlschlegel, Jeff A. Long

## Abstract

Maintaining a stem cell population while developing lateral organs is a crucial aspect of plant development. Studies have shown that a family of micro proteins, the LITTLE ZIPPERS (ZPR), are involved in this process by repressing the activity of HD-ZIP III transcription factors. However, the unique role of each ZPR has not been thoroughly characterized. In this work, we use genetics, imaging, and biochemistry to create a detailed picture of ZPR family expression and their specific interactions with HD-ZIP IIIs. CRISPR/Cas9 was implemented to isolate single loss-of-function ZPR alleles as well as higher-order mutant combinations. A single mutation in *ZPR1, ZPR3*, and *ZPR4* affects the development of the cotyledons during embryogenesis. Additionally, double mutant analyses indicates both unique roles for each ZPR protein as well as redundancy. Using ZPR tagged lines we show that while ZPR3 and ZPR4 have a similar pattern of accumulation as the HD-ZIP IIIs, ZPR1 and ZPR2 accumulation is more limited. Immuno-precipitations (IP) with tagged ZPR proteins are mainly enriched with the anticipated HD-ZIP III partners. Although ZPRs interact with all HD-ZIP IIIs, an apparent preference of heterodimer formation with REVOLUTA is observed. Our work highlights that the dynamics of ZPR protein accumulation together with the strength of ZPR-HD-ZIP III interactions provide an added layer of complexity to the regulation of HD-ZIP IIIs during plant development.

## Introduction

During *Arabidopsis* embryogenesis, the embryo establishes an apical/basal axis. Both the apical domain and the basal domains will develop a population of stem cells. The shoot apical meristem (SAM) will give rise to all the “above-ground” organs, and the root meristem, derived from the basal domain, will give rise to the root system. The SAM consists of a dome of cells which can be broken down into zones and layers based on their function. The central zone of the SAM is dedicated to the maintenance of stem cell identity while the peripheral zone gives rise to new organs (Soyars et al., 2016). The balance between maintenance and proliferation in the SAM is a fundamental process for proper plant development.

SAM maintenance is governed by the conserved CLV-WUS pathway (Mayer et al., 1998; Somssich et al., 2016; Yadav et al., 2011), and the spatial coordination of primordia production (phyllotaxis) is coordinated by polar auxin transport (Bhatia et al., 2016). New leaves develop from the periphery of the meristem and acquire a distinct polarity along the adaxial/abaxial axis. The adaxial domain of the leaf is toward the center of the SAM and will become the upper part of the leaf. The abaxial domain is away from the center of the meristem and will become the lower part. Precise ad/abaxial patterning is very important for SAM formation and maintenance as well as for correct leaf development (Eshed et al., 2001; McConnell & Barton, 1998).

CLASS III HOMEODOMAIN LEUCINE-ZIPPER (HD-ZIP III) transcription factors are major positive regulators of adaxial polarity and play a central role in meristem formation and maintenance. *The Arabidopsis* genome contains five HD-ZIP III genes [*REVOLUTA/IFL1* (*REV*), *PHABULOSA/ATHB14* (*PHB*), *PHAVOLUTA/ATHB9* (*PHV*), *CORONA/ATHB15* (*CNA*) and *ATHB8*]. Members of this group share five predicted structural domains: A Homeodomain - a DNA binding domain, a leucine zipper domain – a protein dimerization domain, a START domain - a motif with putative lipid binding capability (Nagata & Abe, 2021), a SAD domain - a putative transcriptional activation domain (De Caestecker et al., 2000), and a MEKHLA domain – a protein-protein interaction domain, present exclusively in the class III members, possibly involved in oxygen redox and light signaling (Chandler et al., 2007; Duclercq et al., 2011; Magnani & Barton, 2011; Mukherjee & Bürglin, 2006).

HD-ZIP IIIs play essential roles in *Arabidopsis* shoot development. This includes establishment of the embryonic SAM, patterning and polarity of vascular tissues, and establishment of leaf polarity (Emery et al., 2003; Mallory et al., 2004; McConnell et al., 2001; Otsuga et al., 2001; Prigge et al., 2005; Smith & Long, 2010; Talbert et al., 1995). Using loss-of-function (lof) alleles, genetic studies revealed a complex pattern of unique, overlapping and antagonistic functions of HD-ZIP IIIs (Prigge et al., 2005). *rev* is the only single mutant that displays an obvious phenotype on its own. Plants mutant for REV have revolute leaves and display a reduction in axillary meristems (Talbert et al., 1995). The PHB and PHV proteins perform overlapping functions with REV as well as CNA. Addition of a *rev* lof allele to *phb phv* double mutant leads to loss of the SAM and abaxialization of the cotyledons. Addition of *cna* lof allele to *phb phv* has the opposite effect on the SAM, resulting in a larger meristem. This suggests that CNA has functions that are both antagonistic to those of REV within certain tissues and overlapping with REV in other tissues (Prigge et al., 2005).

Each *HD-ZIP III* has a unique pattern of expression and these patterns share a spatial-temporal overlap in different tissues during the development. For example, *PHB/PHV/REV/CNA* are expressed in the apical domain of the embryo at the globular stage, and determine the formation of the SAM and the cotyledons (Emery et al., 2003; Prigge et al., 2005; Smith & Long, 2010). Later in development, these four family members are also expressed in the adaxial regions of the cotyledons and all five *HD-ZIP III*s are expressed in the pro-vasculature of the future hypocotyl and root (Grigg et al., 2009; Lucas et al., 2013; Smith & Long, 2010; Williams et al., 2005). During the post-embryonic growth phase, four *HD-ZIP III*s are expressed in the SAM to maintain its meristematic activity and during the initiation of lateral meristems. They are also expressed in the adaxial domain of newly forming leaf primordia to establish the polarity of these tissues (Emery et al., 2003; McConnell et al., 2001; Prigge et al., 2005).

HD-ZIP III expression, protein accumulation and function are tightly controlled by multiple factors. The expression domain of each of the five *HD-ZIP III* genes is determined by their promoters as well as by cell type-specific miRNA action. *HD-ZIP III*s are post-transcriptionally regulated by the miR165/166 family which bind to the miRNA binding site in HD-ZIP III coding sequence. This triggers the degradation of HD-ZIP III mRNAs, thereby limiting their mRNA accumulation to a subset of cells in which they are transcribed (Byrne, 2006; Juarez et al., 2004; Kidner & Martienssen, 2004; J. Kim et al., 2005; Prigge et al., 2005; Williams et al., 2005; Xie et al., 2005).

HD-ZIP IIIs are also controlled on the post-translational level by the LITTLE ZIPPER (ZPR) proteins. The *Arabidopsis* ZPR family consists of four micro proteins (67-105 amino acids) with a single leucine-zipper (LZ) domain (Sup. Fig. 1A). These micro proteins were suggested to repress HD-ZIP III function by forming dimers with HD-ZIP IIIs through their shared LZ domains, and blocking HD-ZIP III DNA binding. In addition, it was suggested that HD-ZIP IIIs bind to the promoter of ZPR genes and regulate their expression (Y. S. Kim et al., 2008; Wenkel et al., 2007).

ZPR proteins can be divided into two subgroups. All four proteins share a highly homologous leucine zipper domain. ZPR3 and ZPR4 start with the leucine zipper domain, followed by an asparagine and serine rich tail. ZPR1 and ZPR2 sequences are longer and have an additional peptide sequence before and after the LZ domain. A functional role for these additional sequences has not been investigated. Currently, little is known about the expression of the *ZPR* genes and their protein accumulation at multiple stages of development. *ZPR1* and *ZPR3* have been reported to be expressed in the adaxial side of the embryonic cotyledons, as well as in the meristem and the pro-vasculature by *in situ* hybridization (Wenkel et al., 2007). *ZPR2* expression was detected only in the organizing center of the vegetative and reproductive SAM (Weits et al., 2019). *ZPR4* expression has not been described.

ZPR2 and ZPR3 loss-of-function mutant phenotypes were previously described, highlighting the role of these genes in meristem function. A *ZPR3* mutant (*zpr3-2*) had shorter internodes between siliques and in some plants, two or more secondary inflorescences developed in the cauline leaf axils. These observations indicate that the activities of the SAM and/or axillary meristems are disrupted in *zpr3-2* (Y. S. Kim et al., 2008). *ZPR2* mutants (*zpr2-2* and *zpr2-3)* had a lower leaf initiation rate in comparison to wild-type, indicating a role for ZPR2 in maintenance of timing of leaf production by the SAM (Weits et al., 2019). *zpr3 zpr4* double mutant seedlings produced extra cotyledons, larger meristems and more rosette leaves. In addition, development of multiple ectopic meristem-like structures was observed around the primary SAM. This indicates that *ZPR3* and *ZPR4* have a redundancy in their role during embryogenesis, are essential for proper functioning of stem cells in the SAM, and proper development of leaves (Y. S. Kim et al., 2008).

Neither the unique role of each ZPR nor the redundancy between them has been thoroughly studied. Little is known about the difference in their expression patterns throughout development and nothing has been reported about their intracellular localization *in vivo*. It is also not known whether they have a preference in their interaction with specific HD-ZIP IIIs. Furthermore, the dynamics of their interaction with HD-ZIP IIIs and the regulation of these interactions is not clear. In this work, we employ genetics, imaging, and biochemistry to analyze ZPR family accumulation patterns and their specific interactions with HD-ZIP IIIs. Here we provide a characterization of ZPR expression, mutant phenotypes and protein interactions that sheds light on the unique attributes and functions of each ZPR protein.

## Results

### 1. The unique role of each ZPR and the redundancy of their function

To obtain insight into the molecular mechanism behind ZPR action, we aimed to characterize single mutants and combinations of the higher-order mutants. Previous work has described the effect of T-DNA insertions in several *Arabidopsis ZPR* genes (Y. S. Kim et al., 2008; Weits et al., 2019; Wenkel et al., 2007). In this work we have implemented CRISPR/Cas9 to create lof alleles for all four *ZPR* genes. For each *ZPR* we have isolated a lof allele with mutations in the coding sequence resulting in loss of the LZ domain (Sup. Fig. 2).

A *zpr1* mutant (*zpr1-2*, Sup. Fig. 2) produces seedlings with three cotyledons (tricots) as well as two fused cotyledons (monocots) at a higher frequency in comparison to Ler-0 (Fig. 1A, B, N), indicating a unique role for ZPR1 during embryogenesis. A *zpr2* mutant (*zpr2-4*, Sup. Fig. 2) was indistinguishable from Ler-0 and no tricot seedings were observed (Fig. 1C, N). Previously it was shown that *zpr2* mutant plants had a lower leaf initiation rate; however, this phenotype was not observed in *zpr2-4* under our growth conditions (Weits et al., 2019). The difference between these *zpr2* alleles might be attributed to the different ecotypes used in these studies. Similar to *zpr1-2*, *zpr3* mutants (*zpr3-3*, Sup. Fig. 2) also had a higher ratio of tricot formation (Fig. 1D, N). The phyllotaxy and internode height of the cauline leaves in *zpr3-3* mutant were affected in a similar manner as a previously described *zpr3-2* allele (Y. S. Kim et al., 2008) (Sup. Fig. 3A, B). In addition, a higher number of carpels was observed in the *zpr3-3* mutant (Fig. 1F, H, I, K, L, O), resembling the phenotype of lof *hd-zip III* mutants (Prigge et al., 2005). A *zpr4* single lof mutant was characterized in previous work (Y. S. Kim et al., 2008), which stated that it was indistinguishable from wild-type. However, here we report that a *zpr4* mutant (*zpr4-3*, Sup. Fig. 2), had a higher occurrence of seedlings with three cotyledons (Fig. 1E, N). Taken together, the phenotype of the single lof *zpr* mutants suggests a unique role for each ZPR protein and highlights their importance in embryogenesis, meristem function and organ initiation.

**Figure 1.**
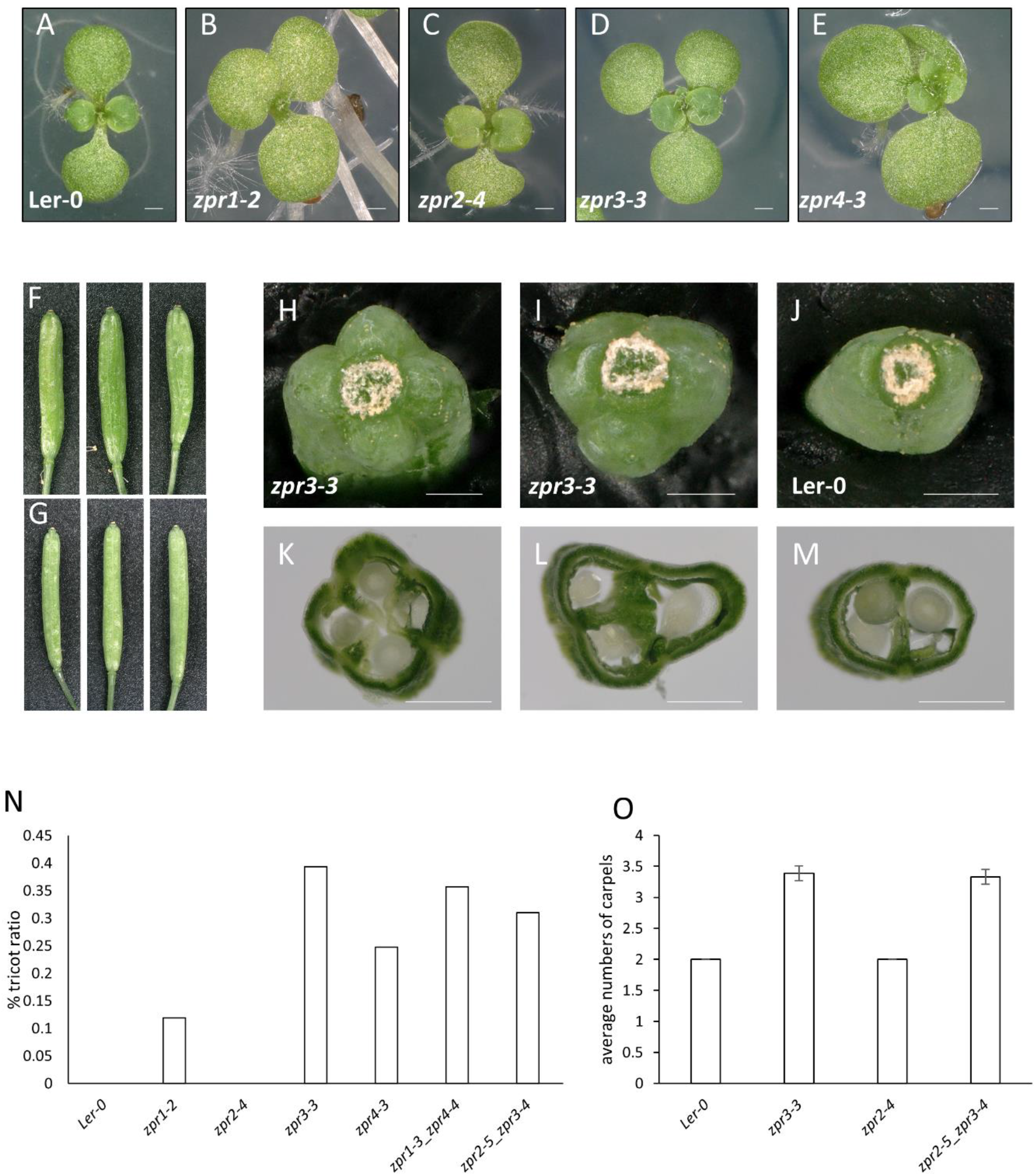
Phenotypes of loss of function ZPR alleles created by CRISPR/Cas9. a-e, 10 days old ZPR mutant seedlings with 2 or 3 cotyledons in comparison to the Ler-0 seedling with 2 cotyledons. f-m, Representative siliques of zpr3-3 mutant with 3 or 4 carpels (f,h,i,k,l) in comparison to Ler-0 siliques with 2 carpels (g,j,m). Scale bar - 500 μm. n, ratio of penetrance in *zpr* mutants for seedlings with three cotyledons in comparison to Ler-0. Number of plants used for each line is: 673 Ler-0, 840 *zpr1-2*, 1015 *zpr2-4*, 254 *zpr3-3*, 404 *zpr4-3*, 561 *zpr1-3_zpr4-4*, 323 *zpr2-5_zpr3-4.* o, Average numbers of carpels in siliques 1 through 10 from 10 ZPR mutant plant lines in comparison to Ler-0. Error bars indicate ±2*SE of the mean for 100 individual siliques.

Next, we created double mutants to explore the redundancy of *ZPR* genes. Using Col-0 T-DNA alleles, a *zpr3-2 zpr4-2* double mutant was previously characterized (Y. S. Kim et al., 2008). These plants formed extra cotyledons, had an enlarged meristem and formed ectopic SAM-like structures that led to a bushy appearance with multiple shoots. Using newly generated alleles in the Ler-0 background, we characterized a *zpr3-5 zpr4-5* double mutant and saw many of the same phenotypes (Sup. Fig. 2, Sup. Fig. 3 C-E). Flowers from the *zpr3-5 zpr4-5* were severely affected and produced extra floral organs and unfused gynoecia (Sup. Fig. 3E). These plants were unable to produce any seeds. These severe phenotypes suggest ZPR3 and ZPR4 play redundant roles regulating organ formation and SAM function throughout the life of the plant in *Arabidopsis*.

To determine the extent of possible redundancy between other ZPR family members, we have also created and characterized two additional pairs of double mutants, *zpr1-3 zpr4-4* and *zpr2-5 zpr3-4*. These combinations did not result in an enhancement of either the tricot phenotypes seen in *zpr1, zpr3, and zpr4* single mutants or the carpel number phenotype seen in the *zpr3* single mutants (Fig. 1N, O). This suggests that ZPR1 is not redundant with ZPR4 and ZPR2 is not redundant with ZPR3, or that their redundancies are masked by the function of another ZPR family member. Additional combinations of ZPR mutants might shed light on the redundancy of roles of ZPR proteins.

### 2. ZPR expression and cellular localization in different stages of development

To characterize the accumulation pattern of ZPR proteins we used recombineering to create plants expressing ZPR - fluorescent protein fusions within large chromosomal contexts (Alonso & Stepanova, 2014; Brumos et al., 2020) (Fig. 2A). Using these ZPR tagged lines we have detected ZPR1, ZPR3 and ZPR4 in the metaxylem of the root – a known domain of HD-ZIP III expression (Carlsbecker et al., 2010; Ramachandran et al., 2016), while ZPR2 was not detected in this tissue (Fig. 2B-E). This suggests a role for ZPR regulation of HD-ZIP IIIs in development of vascular tissues, although we did not observe any obvious root growth phenotypes in our single and double lof combinations.

**Figure 2.**
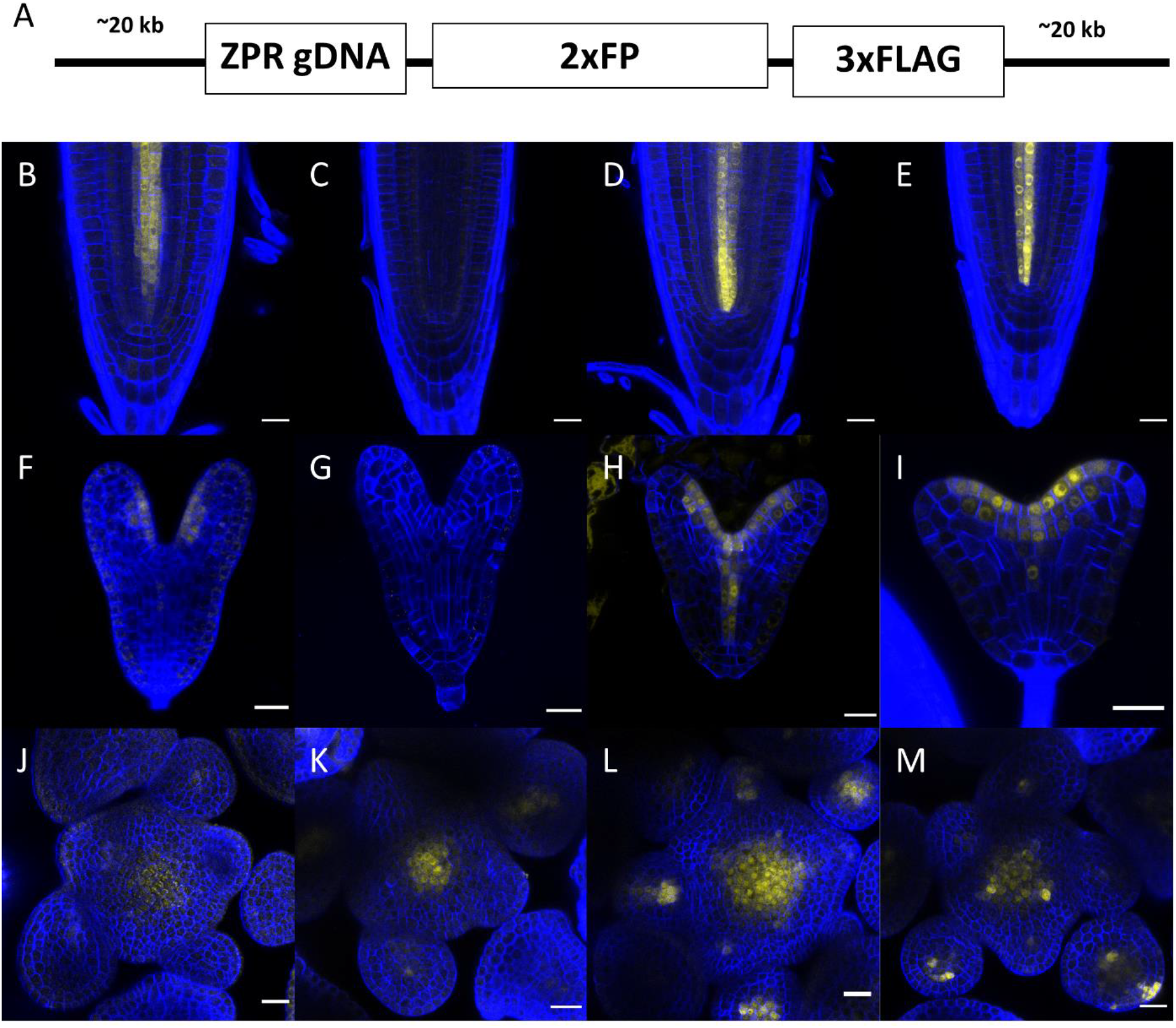
ZPRs expression pattern in different tissues. a. Scheme of a C-terminal ZPR-tagged recombineering construct. ZPR genomic DNA fused to 2 x Florescent Protein and 3 x FLAG tag. Accumulation of ZPR proteins - ZPR1 (b,f,j), ZPR2 (c,g,k), ZPR3 (d,h,l), ZPR4 (e,I,m) in seedling roots 5 days after germination (b-e), in embryos (f-i), in the inflorescence (j-k). Scale bar = 20 μm.

In a previous report describing *ZPR1* and *ZPR3* expression in the embryo, it was noted that the expression of these *ZPR* genes could not be detected earlier than the heart stage of embryogenesis (Wenkel et al., 2007). However, we detected ZPR3 protein in the apical domain of the early globular stage embryo (Sup. Fig. 4). At the heart stage, ZPR3 expression resembles HD-ZIP III expression - the protein is found in the meristem, in the vasculature, and in the adaxial side of the cotyledons (Fig. 2H). ZPR4 accumulation was very similar to ZPR3 throughout embryo development (Fig. 2I and Sup. Fig. 5), though ZPR3 appears to be slightly more abundant, especially in the SAM. ZPR3 and ZPR4 accumulation in the early stages of embryogenesis resembles the spatial-temporal expression of *REV, PHB, PHV and CNA* at these stages (Emery et al., 2003; McConnell et al., 2001; Otsuga et al., 2001; Smith & Long, 2010).

While ZPR3 and ZPR4 accumulation is very similar throughout embryo development and resembles HD-ZIP III expression, ZPR1 can be detected only in the later stages of embryogenesis (in the heart and torpedo stage). In addition, ZPR1 accumulation levels are lower in the adaxial side of the cotyledons and in the vasculature in comparison to ZPR3/4, and is barely detected in the embryonic meristem (Fig. 2F). A similar expression pattern was also seen by *in-situ* hybridization of *ZPR1* (Wenkel et al., 2007). ZPR2 accumulation was not detected at any stage of embryogenesis (Fig. 2G). ZPRs expression pattern in the adaxial side of the cotyledons, in the vasculature and in the area that will later become a meristem, highlights the essential role of ZPR3 and ZPR4 in embryo development, while ZPR1’s role is narrower and ZPR2 may not have a role during embryogenesis.

Though ZPR2 was not detected in the root vasculature and during embryogenesis, we were able to detect ZPR2 protein accumulation in the inflorescence meristem (Fig. 2K), similarly to what has previously been reported in the vegetative SAM (Weits et al., 2019). ZPR3 and ZPR4 accumulation in the inflorescence resembled ZPR2, but was broader and more readily detectable (Fig. 2L, M). *In situ* hybridization assays revealed that ZPR3 expression is confined to the CZ with a higher density in the organizing center, where WUS is also highly expressed (Y. S. Kim et al., 2008). ZPR1 was detected in the inflorescence meristem albeit its accumulation was very low (Fig. 2J).

While HD-ZIP III proteins are mainly localized to nuclei (Caggiano et al., 2017; Carlsbecker et al., 2010; Donner et al., 2009; Smith & Long, 2010), ZPR2, ZPR3 and ZPR4 proteins are distributed between the nucleus and the cytosol and ZPR1 was mainly found in the cytosol (Fig. 2B, D, E). The observation of ZPR3 cytoplasmic expression was previously noted. When expressed transiently, ZPR3 could be found in the nucleus as well as in the cytoplasm (Husbands et al., 2016; Y. S. Kim et al., 2008). The presence of ZPRs in different cellular compartments suggests a regulatory mechanism of their trafficking into the nucleus that might affect their interaction with HD-ZIP IIIs. Another thing that should be noted is the dynamic nature of ZPR3 and ZPR4 expression in different cells (Sup. Fig. 4 and 5). The proteins are not evenly distributed in all expressing cells, with varying levels of accumulation in different cells and tissues. In addition, in some cells the ZPR protein seems to accumulate more in the nucleus, while in other cells it is more present in the cytosol.

### 3. ZPR protein-protein interaction partners

Transcription factors often must interact with other proteins to function (Amoutzias et al., 2008; Riechmann et al., 2000). HD-ZIP IIIs were shown *in vitro* to form dimers through their LZ domain in order to bind to DNA (Sessa et al., 1998). ZPRs were hypothesized to competitively inhibit HD-ZIP III function by creating non-functional heterodimers through their shared LZ motifs, thereby preventing HD-ZIP III dimerization and binding to their target genes (Y. S. Kim et al., 2008; Wenkel et al., 2007).

The interactions between *Arabidopsis* HD-ZIP III and ZPR proteins have been evaluated through yeast two-hybrid and *in vitro* pull-down assays. ZPR3 was shown to interact with all five HD-ZIP III proteins (Y. S. Kim et al., 2008) and REV was shown to interact with all four ZPR proteins (Wenkel et al., 2007). Overlap of ZPR expression patterns within the family, as well as their overlap with HD-ZIP III expression, raises questions about the combinatorial nature of interactions between these families on the protein level.

To explore ZPR interactors *in planta* we used flowers expressing 3xFLAG tagged ZPR proteins to perform an immunoprecipitation and MS analysis. Anti-FLAG blots of the elution fraction (Fig. 3A) show enrichment of the tagged ZPR protein in the experiment samples in comparison to the Ler-0 control. MS/MS analysis revealed that the most highly enriched proteins in the immunoprecipitated fraction were the HD-ZIP IIIs. Each ZPR protein interacted with all members of the HD-ZIP III family. In each ZPR IPMS experiment the bait ZPR protein was detected (Fig. 3C), but the absence of other ZPR proteins from the pulled down fraction might suggest that ZPRs do not form heterodimers with other ZPRs. Taken together, this suggests that HD-ZIP IIIs are the only proteins with which ZPRs interact through their LZ motif. No other LZ-containing proteins were found to interact with ZPRs in our experiments.

**Figure 3.**
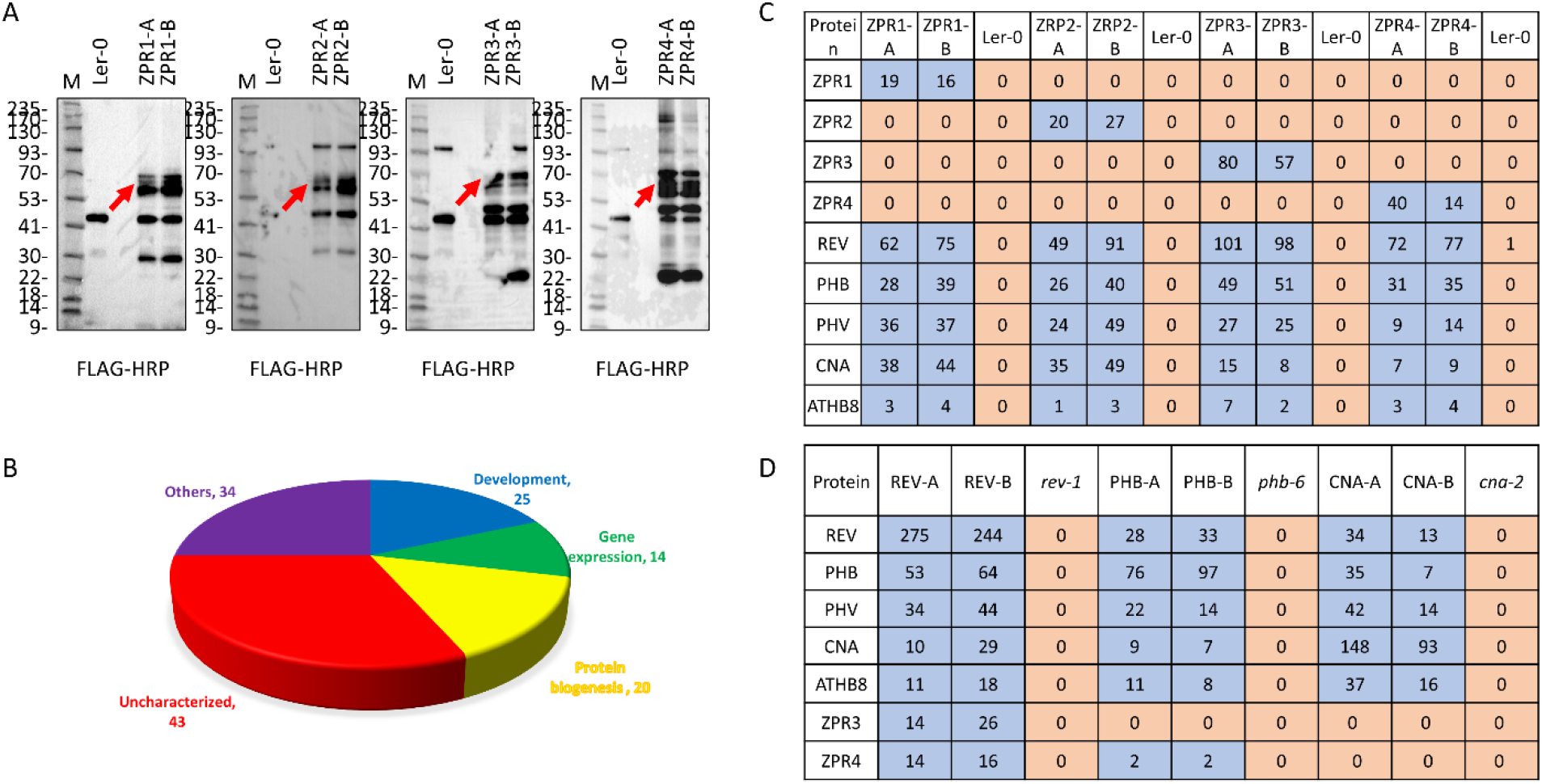
ZPR protein-protein interactions. a, An anti-FLAG blots of the elution fraction. Red arrows point to an enrichment of the tagged ZPR protein in the two independent biological replicate lines in comparison to the Ler-0 control. b, Pie chart showing PANTHER GO annotation and classification of ZPR interacting partners. c, ZPR protein-protein interactions. d, HD-ZIP III protein-protein interactions.

An apparent ZPR hierarchy of preference for specific HD-ZIP IIIs was observed in these experiments. Higher MS/MS counts of REV were measured in all ZPR IP samples followed by PHB, PHV and CNA and much lower counts of ATHB8 (Fig. 3C). A strong REV-ZPR interaction was also observed in the reciprocal IPMS experiment using 3xFLAG tagged HD-ZIP III proteins (Sup. Fig. 6). REV pulled down ZPR3 and ZPR4, while ZPR4 was also pulled down with PHB, and no ZPRs were detected in CNA immunoprecipitation (Fig. 3D). This could suggest that ZPRs have a higher affinity for interactions with REV.

Immunoprecipitation fractions for each of the HD-ZIP III proteins were mainly enriched with all the HD-ZIP III family members (Fig. 3D). This is the first report of HD-ZIP IIIs heterodimer formation *in planta*, supporting previous *in vitro* observations (Sessa et al., 1998). The highest MS/MS count captured in each experiment was of the bait protein. The MS/MS count is consistent with the idea that more heterodimers were formed between REV and PHB or PHV and less heterodimers were formed between REV and CNA or ATHB8. Similarly, PHB formed more heterodimers with REV and PHV, and less with CNA and ATHB8. Given the semi-quantitative nature of these analyses, however, these hypotheses will require validation using more quantitative means. Surprisingly, the CNA IP fraction contained only the HD-ZIP III proteins and no other protein was pulled down with CNA. CNA had a similar MS/MS count for the other four HD-ZIP IIIs, suggesting a similar amount of heterodimer formation between CNA and REV/PHB/PHV/ATHB8. This divergence in MS/MS count sheds light on the preferred pattern of hetero-dimer formation between different HD-ZIP IIIs and supports the genetic interactions previously observed (Prigge et al., 2005).

PANTHER GO anthology (Thomas et al., 2003) was used to annotate and classify the majority of the one hundred thirty-six ZPR pulled down proteins (Fig. 3B). The pull-down fractions were enriched with twenty-five proteins involved in different aspects of development (Sup. Table 1). Due to the presence of HD-ZIP IIIs and ZPRs genes in the input list, biological processes such as determination of bilateral symmetry, adaxial/abaxial pattern specification, meristem development, axis specification and embryo development were highly enriched and included the following proteins: LUG, FIP37, VCS, RNJ (EMB2746), FAB1 (KAS2), CRU3 (CRC), emb2386, SPP (AT5G42390) and EDD1.

Developmentally related proteins had additional GO classifications such as gene expression, translation, protein metabolism, protein modification and transport. Searching for these GO classifications upon the pulled down proteins resulted in classification of thirty-four additional proteins - fourteen involved in gene expression and chromatin organization (Sup. Table 2) and twenty involved in protein biogenesis (Sup. Table 3). Another group of forty-three proteins were classified as unmapped or unclassified (Sup. Table 4), suggesting there is a great potential of revealing new and previously uncharacterized molecular mechanisms involving ZPRs. The residual thirty-four proteins were grouped as others (Sup. Table 5).

The ZPR1 and ZPR2 IP fractions were enriched with proteins involved in protein proteo-stasis: protein synthesis (ribosomal proteins), protein degradation (proteases and ubiquitin protein ligase), post-translational modifications and transport machineries (Sup. Table 3). This correlates nicely with detection of ZPR proteins in the cytosol as well as in nuclei and suggests a post-translational regulation mechanism affecting ZPR cellular dynamics, protein interactions, or other aspects of ZPR function. Post-translational modification of ZPR2 was shown to be an important check point for ZPR2 protein accumulation in the SAM stem cell niche (Weits et al., 2019). ZPR1 also has the N-terminal Met-Cys motif and might be part of this N-degron regulation. External signals and intrinsic developmental cues may affect the conformation of the HD-ZIP III and ZPR proteins through post-translational modifications, allowing them to recognize each other and form heterodimers.

DNA binding proteins were also detected between ZPR1 and ZPR2 interacting proteins (Sup. Table 2). For example, NAP1;3 (AT5G56950.1) was pulled down with ZPR1 and ZPR2, while FKBP53 (AT4G25340.1) and VIP3 (AT4G29830.1) were pulled down with ZPR2. Additional nucleosome assembly proteins such as PRMT4B (AT3G06930.2) and AT5G49020.1 were pulled down with ZPR1, and AT3G10690.1, AT2G30620.1 and AT5G65360.1 were pulled down with ZPR2. This might suggest that ZPRs do associate with DNA independently of HD-ZIP IIIs, possibly through the peptide extensions unique to ZPR1 and ZPR2.

## Discussion

ZPRs are micro proteins that are thought to have originated as a duplication of the HD-ZIP III leucine zipper domain. They represent an additional regulatory model on HD-ZIP III and contribute to the complexity of HD-ZIP III activity (Floyd et al., 2014). ZPR expression levels are regulated by the HD-ZIP IIIs, and they form dimers (Y. S. Kim et al., 2008; Wenkel et al., 2007) or tetramers (Husbands et al., 2016) with HD-ZIP IIIs to fine-tune their function, thus post-translationally moderating the effects of the expressed HD-ZIP IIIs.

HD-ZIP III precise spatial-temporal expression pattern is very important for their function and therefore tightly regulated by different transcription factors [KANADIs, YABBYs, PLT and HD-ZIP IIIs (Smith & Long, 2010)] and by miRNA165/166 activity (Zhou et al., 2007; Zhu et al., 2011). If ZPRs are expressed in the same domain, bind to HD-ZIP IIIs and prevent their binding to DNA, then how do HD-ZIP IIIs execute their function? This suggests that there should be a mechanism facilitating ZPR - HD-ZIP III interaction and separation, allowing them to co-exist. HD-ZIP III - ZPR interaction might depend on their expression levels and patterns of protein accumulation, or on the affinity of the LZ sequences.

Detection of all the members of the HD-ZIP III family in all the ZPR IP experiments suggests that all LZ motifs of different ZPRs are compatible to interact with all the LZ motifs of different HD-ZIP IIIs. Furthermore, all the HD-ZIP IIIs were able to form heterodimers within the family. A mutagenesis analysis where individual Leu and Ile residues were mutated to Ala showed that Leu residues in heptad two, four and six (but not Ile in heptad three) are crucial for the ZPR3-REV (DTM-SlREV) interaction in tomato (Xu et al., 2019). Since all the ZPRs and all the HD-ZIP IIIs have identical residues necessary for LZ interaction (Leu/Ile in the *d* position as well as Asn in *a* position of the heptad) (Sup. Fig. 1B), we can conclude that the specificity of ZPR – HD-ZIP III interaction in *Arabidopsis* is not based on the core of the LZ motif sequence. Although the compatibility with other LZ proteins might be based on those residues and explain the lack of any other LZ domain containing proteins in HD-ZIP III and ZPR IPMS.

While positions *a* and *d* are important for the hydrophobic interaction between the two alpha helices, the dimerization specificity is primarily determined by polar interactions involving the side chains of residues at the *e/g* positions. In addition, the amino acids outside the leucine heptad repeats might determine interaction potential due to possible structural interferences (Deppmann et al., 2004). A minor difference between ZPR3/4 or ZPR1/2 can be seen in the polar side chains of amino acids (Sup. Fig. 1B, black asterisk). For example, the difference between ZPR3 and ZPR4 can be explained by E/Q residues and in other cases the difference between ZPR1 and ZPR2 can be explained by I/R, K/A K/N residues (Sup. Fig. 1B blue asterisk).

In light of the strong interaction between ZPRs and HD-ZIP IIIs, it was puzzling why we could not detect ZPRs forming heterodimers with other ZPRs. We can hypothesize that ZPR dimerization is restricted by a 3D conformation, preventing the LZ domain from being readily available for interaction. Hence ZPR dimerization with HD-ZIP III might rely on a post translational modification mechanism, affecting the protein confirmation, promoting its trafficking into the nucleus and facilitating the interaction with HD-ZIP III. Potential candidates involved in the ZPR post translational regulation mechanism facilitating ZPR - HD-ZIP III interaction were detected in the IP fraction. Moreover, more interacting proteins were detected in ZPR1/2 IP fraction, indicating a possible role for the additional peptide domain found in these proteins.

Differences in interaction strength have not been previously reported for *Arabidopsis* ZPR and HD-ZIP III pairs, nor has the hetero-dimerization within each family have been demonstrated *in planta*. Our work demonstrates the ability of all HD-ZIP IIIs to form heterodimers within the family and form heterodimers with all the ZPRs. IPMS results also show the difference in the strength of different heterodimeric connections. All ZPR proteins appeared to have a higher affinity for REV. Further supporting the stronger interaction between REV and ZPRs, some *zpr3-1d* mutants displayed bare cauline leaf axils lacking lateral meristems (Y. S. Kim et al., 2008) - a phenotype similar to the *rev-1* mutant (Talbert et al., 1995). This suggests that ZPR3 is able to bind specifically to REV in the domain where HD-ZIP III activity is uniquely executed by REV. In addition, DTM - a tomato ZPR3 homolog - was shown to interact with several members of the tomato HD-ZIP III family, but had the highest interaction with REV, suggesting that DTM controls SAM development mainly through suppression of SlREV activity (Xu et al., 2019), and highlighting the conserved role of REV-ZPR protein-protein regulation. The differences in the ability of unique ZPR proteins to interact with different HD-ZIP III proteins, and possible differences in interaction strength, provide an added layer of complexity in HD-ZIP III regulation.

The accumulation pattern of each ZPR protein correlates nicely with the phenotype of lof single and double mutants and reveals the redundancy, as well as the possible specificity of their function. Combining the expression domain with the development variations of the mutants suggests that ZPR1, ZPR3 and ZPR4 have an important role during embryogenesis in apical domain patterning and development of the cotyledons. ZPR2 together with ZPR3 and ZPR4 are important for accurate meristem function and organ initiation (Kim et al., 2008; Weits et al., 2019). Higher expression levels of ZPR3, and the stronger phenotype of new lof alleles in comparison to other ZPR mutants, highlight the importance of this gene in embryogenesis, meristem function and silique development. ZPR2, which is specifically expressed in the organizing center, was also shown to be a sensor for the low oxygen level in the meristem, hence connecting external signals with meristem activity and organ development (Weits et al., 2019). ZPR roles in different stages of development, especially for meristem maintenance and function, organ initiation and phyllotaxy, and embryonic apical domain formation were previously described in different species (Damodaran et al., 2019; Y. S. Kim et al., 2008; Weits et al., 2019; Wenkel et al., 2007; Xu et al., 2019). Together these results suggest that a complicated balance of HD-ZIP III/ZPR interactions regulate patterning of the shoot system throughout development.

## Methods

### Growth Conditions

Plants were grown on soil or Petri plates containing Linsmaier and Skoog (LS) salts medium and 0.8% Agar. Prior to seeds sowing on soil, they were stratified in 0.1% sterile agarose for two-three days at 4°C. Prior to seeds plating on plates, they were surface sterilized with chlorine gas vapor-phase sterilization (Lindsey et al., 2017) and stratified for two-three days at 4°C. All plants were grown at 22 to 24°C under 16-hour light / 8-hour dark cycle conditions. Landsberg (Ler-0) wild-type plants were used to create CRISPR/CAS9 mutant lines and reporter constructs. All transgenic plants were obtained by *Agrobacterium tumefaciens*-mediated floral dip method (Clough & Bent, 1998). T1 seeds resistant to BASTA or Hygromycin were transplanted to soil.

### CRISPR/CAS9 mutagenesis

CRISPR/Cas9 mutants were generated using a previously published protocol (Yan et al., 2015). Two or four guiding-RNAs were designed for each ZPR gene using http://www.genome.arizona.edu/crispr/ platform (primers are summarized in Sup. Table 6). Vectors containing two or three gRNAs for one or two ZPR genes were used to generate single and double mutants. To enhance Cas9 activity and increase the likelihood of selecting heritable lines, T1 seeds were grown on Hygromycin plates at 28 °C for two weeks. Next the seedlings were transplanted to soil and after a week of acclimation subjected to four rounds of heat-shock at 37 °C for 32 hours and returned to 22 °C for 40 hours (Schindele & Puchta, 2020). The genotypes of T2 plants were confirmed by PCR and sequencing. The Cas9 absence was verified in the T3 plants.

All the mutations occurred in the coding region and resulted in loss of the LZ domain as described in Sup. Fig. 2. *zpr1-2* has a 93 bp deletion in the coding region resulting in loss of the LZ domain. *zpr2-4* has a “c” insertion in the coding region causing a translational frame shift and loss of LZ domain, followed by mismatch. *zpr3-3* has an “a” deletion and several single bp changes, causing a translational frameshift and loss of LZ domain. *zpr4-3* has an 18 bp deletion in the coding region, causing loss of six a.a. in the LZ domain. *zpr1-3 zpr4-4* double mutant has a “g” insertion in *ZPR1* and a one hundred sixteen bp deletion in *ZPR4*. *zpr2-5 zpr3-1* has a “c” insertion in gRNA1 and a “t” insertion in gRNA2 in *ZPR2* and an “a” insertion in *ZPR3*. *zpr3-5 zpr4-5* has a “t” insertion and “ag” deletion in *ZPR3* and a “c” insertion in gRNA1 and an “a” insertion in gRNA3 in *ZPR4*.

### Recombineering

To characterize the accumulation pattern of ZPR proteins and to avoid potential misexpression artifacts, we used recombineering to create plants expressing ZPR - fluorescent protein fusions in the native chromosomal contexts (Alonso & Stepanova, 2014; Brumos et al., 2020). Recombineering constructs were generated as previously described (Warming et al., 2005) with the following modifications. For IP-MS experiments, tdTomato-3xFLAG tag constructs were fused in frame with the C-terminus of ZPR1 (K15kQ7), ZPR2 (JAtY64K12), ZPR3 (JAtY77L19), ZPR4 (JAtY64I06) in a transformation-competent artificial chromosome clone using a bacterial recombineering approach (Alonso & Stepanova, 2014; Brumos et al., 2020). Each construct was transformed via the *Agrobacterium* floral dip method (Clough & Bent, 1998) into Ler-0. Additional lines were similarly generated with 2xYPET-3xFLAG fused to the C-terminus of ZPRs. Primers used for recombineering are described in Sup. Table 7.

### Microscopy

For each ZPR reporter line, two independent transgenic lines were imaged. For tdTomato and YPET imaging, ovules, seedlings and inflorescence meristems were fixed in 4% paraformaldehyde, vacuum infiltrated, rinsed with phosphate buffer saline (PBS), vacuum infiltrated with 2% SCRI Renaissance 2200 (Renaissance Chemicals) and 4% DMSO, then washed twice in PBS. Embryos, seedlings roots and inflorescence meristems were imaged using a Zeiss LSM 710 laser scanning confocal microscope. SR2200 was excited with the ultraviolet diode 405 nm line, and emission was measured between 420 and 470 nm. tdTomato was excited with a 561 nm laser line and emission was measured between 567 and 613 nm. YPET was excited with a 517 nm argon laser line and emission was measured between 500 and 535 nm.

### IP-MS

Immunoprecipitation and mass spectrometry analysis were done as previously described (Moissiard et al., 2014) with some variations. Briefly, 15 grams of inflorescence tissue were ground in liquid nitrogen using a RETSCH homogenizer and resuspended in 25 mL ice-cold IP buffer [50 mM Tris**·**HCl pH 8.0, 150 mM NaCl, 5 mM MgCl2, 0.1% Nonidet P-40, 10% (vol/vol) glycerol, 1x Protease Inhibitor Mixture (Roche)]. Resuspended tissue was homogenized twice using a glass douncer, filtered once through Miracloth and centrifuged 10 min at 16,000 g at 4 °C. The supernatant was diluted to 150 ml. 500 μL M2 magnetic FLAG-beads (SIGMA, M8823) were added to the supernatant and incubated for 120 min rotating at 4 °C. M2 magnetic FLAG-beads were washed five times in ice-cold IP buffer for 5 min rotating at 4 °C. Immunoprecipitated proteins were eluted two times with 600 μL 250ug/mL 3xFLAG peptides (SIGMA, F4799) in TBS [50mM Tris-HCl pH 7.4, 150mM NaCl] for 10 min rotating at RT. Ler-0 wild-type tissue was used as a negative control. IPs were then digested by the sequential addition of lys-C and trypsin proteases, fractionated online using a 25 cm long, 75 μm ID fused-silica capillary that was packed in-house with bulk ReproSil-Pur 120 C18-AQ particles as described elsewhere (Jami-Alahmadi et al., 2021), and then analyzed by tandem mass spectrometry on an Orbitrap Fusion™ Lumos™ Tribrid™ Mass Spectrometer (Thermo Fisher Scientific) with an MS1 resolution of 120,000 followed by sequential MS2 scans at a resolution of 15,000. Raw data were searched against the TAIR *Arabidopsis* reference proteome. Label-free quantitation (LFQ) intensities were calculated by applying the default settings for LFQ analysis using MaxQuant software as described previously by (Cox & Mann, 2008). Proteins that were enriched in two independent biological replicate lines in comparison to the control line of plants were considered true interactors. Overall, seventy-one proteins were detected as ZPR1 interactors, fifty-nine proteins were interacting with ZPR2, fourteen proteins with ZPR3, and eleven proteins with ZPR4.

## Acknowledgements

This work was supported by NIH grant R01GM120623 to J.A.L..

## Author contributions

A.V.G. and J.A.L. designed the experiments. A.V.G. and M.K. generated CRISPR/Cas9 lines. A.V.G. generated reporter constructs, and performed all experiments. A.V.G., Y.J.A. and J.A.W. performed IPMS experiments and analyses. A.V.G. and J.A.L. wrote the manuscript. All authors read and approved the manuscript.

The authors declare no competing financial interests.

---

**Sup. Fig. 1.**
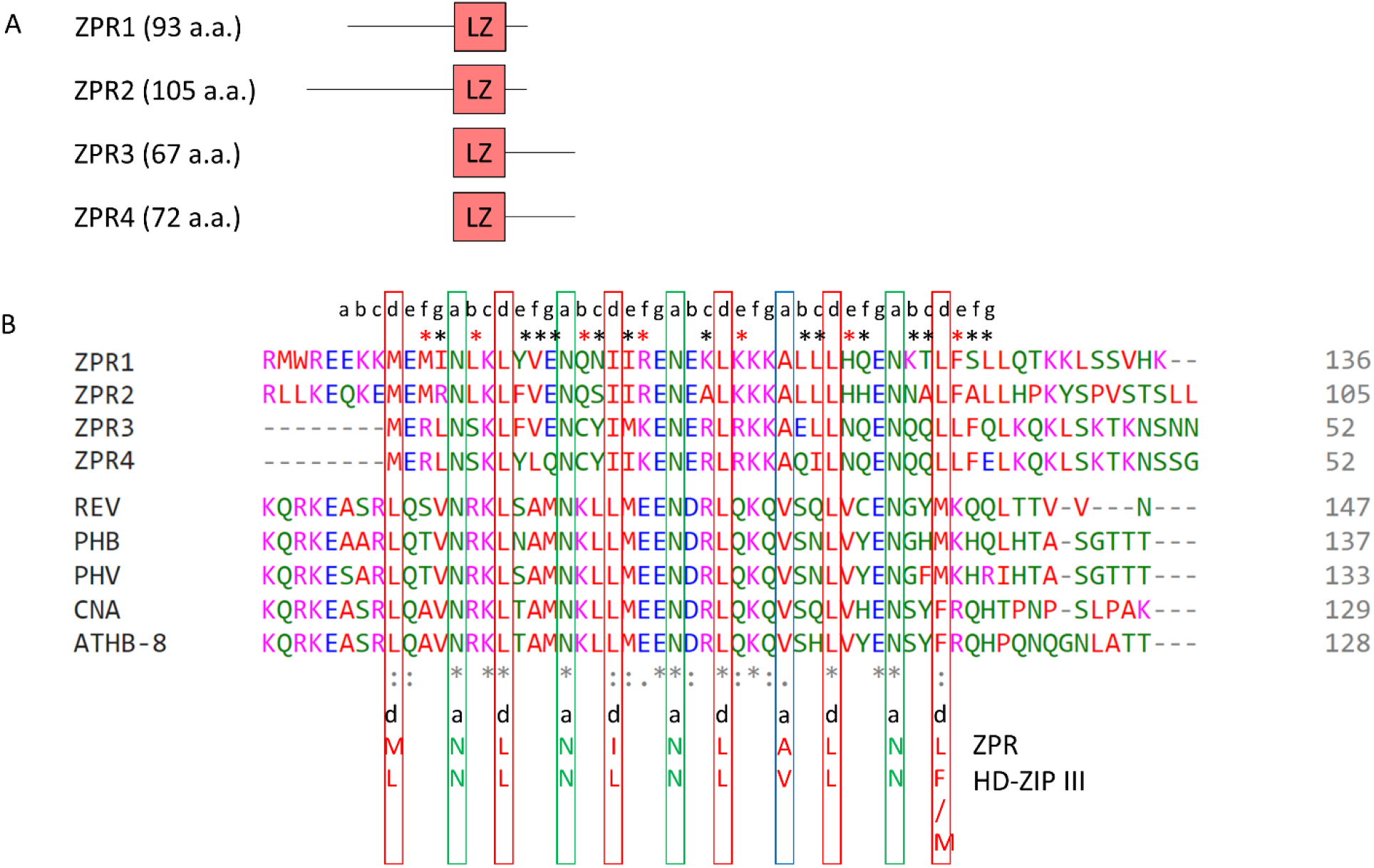
ZPR proteins structure and sequence. Protein domains of *Arabidopsis* ZPR proteins, with the length of each peptide and the leucine zipper (LZ) domain position highlighted in the peptide sequence. b, LZ motif of ZPR proteins aligned with the LZ motif of HD-ZIP IIIs. 6 LZ heptads are shown. Conserved amino acids in positions “a” and “d” are highlighted. Amino acids that differ between the various ZPR homologs are highlighted by black asterisk. ZPR1/2 residues that are different from ZPR3/4 are highlighted by red asterisk.

**Sup. Fig. 2.**
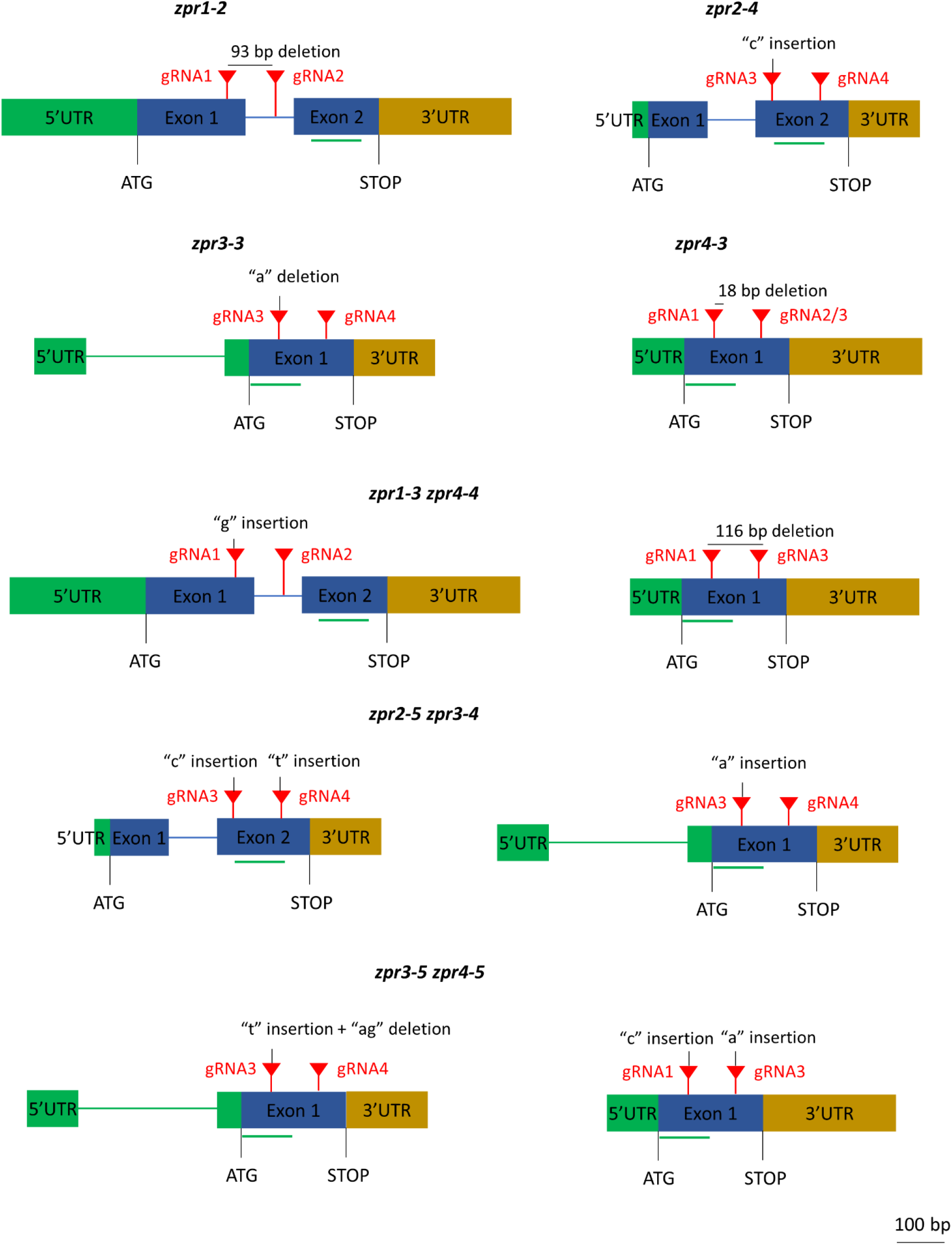
ZPR single and double mutant alleles generated with CRISPR/Cas9. Schematic illustration of the gRNA target design aiming for deletions in the coding region of ZPR proteins. The two single gRNA target sites are indicated by two red triangles. The green line indicates the LZ domain. The deletion or insertion mutations are indicated for each isolated mutant. All the mutations accrued in the coding region and resulted in loss of LZ domain. *zpr1-2* has a 93 bp deletion. *zpr2-4* has a “c” insertion followed by mismatch. *zpr3-3* has an “a” insertion and several single bp changes. *zpr4-3* has a 18 bp deletion. *zpr1-3 zpr4-4* has a “g” insertion in the *ZPR1* and a 116 bp deletion in the *ZPR4*. *zpr2-5 zpr3-4* has a “c” insertion in gRNA1 and a “t” insertion in gRNA2 in the *ZPR2*, and an “a” insertion in *ZPR3*. *zpr3-5 zpr4-5* has a “t” insertion and “ag” deletion in *ZPR3*, and a “c” insertion in gRNA1 and an “a” insertion in gRNA3 in *ZPR4*.

**Sup. Fig. 3.**
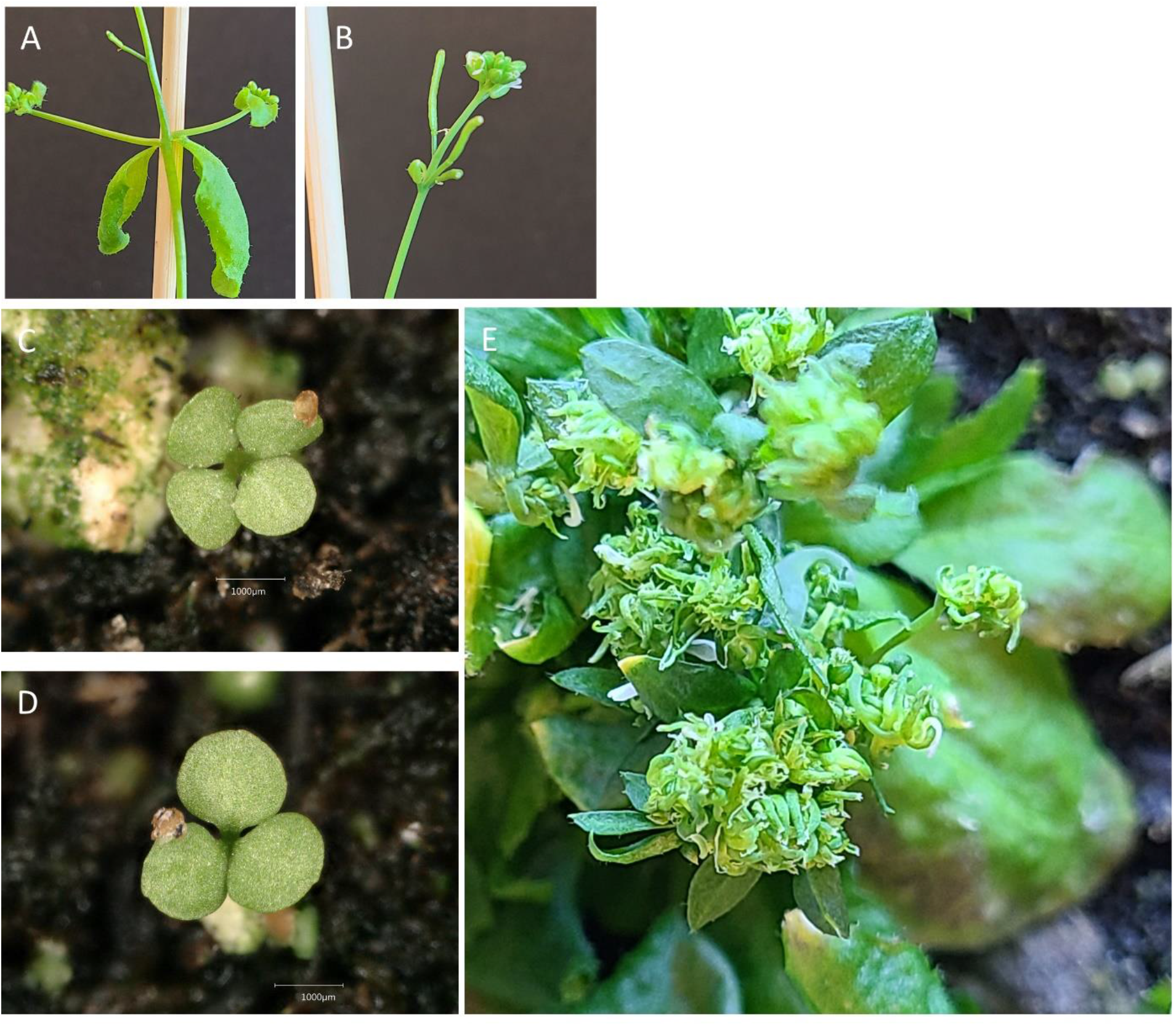
Phenotypes of loss of function ZPR alleles created by CRISPR/Cas9. a-b, representative images of *zpr3-3* mutant. Phenotype of a four weeks old *zpr3-3* mutant plant with shorter internodes between axillary inflorescence (a) and between siliques (b). c-e, representative images of *zpr3-5 zpr4-5* double mutant. 7 days old *zpr3-5 zpr4-5* seedlings with 4 cotyledons (c) and 3 cotyledons (d). 8 weeks old *zpr3-5 zpr4-5* plant with bushy appearance and abnormal inflorescence (e).

**Sup. Fig. 4.**
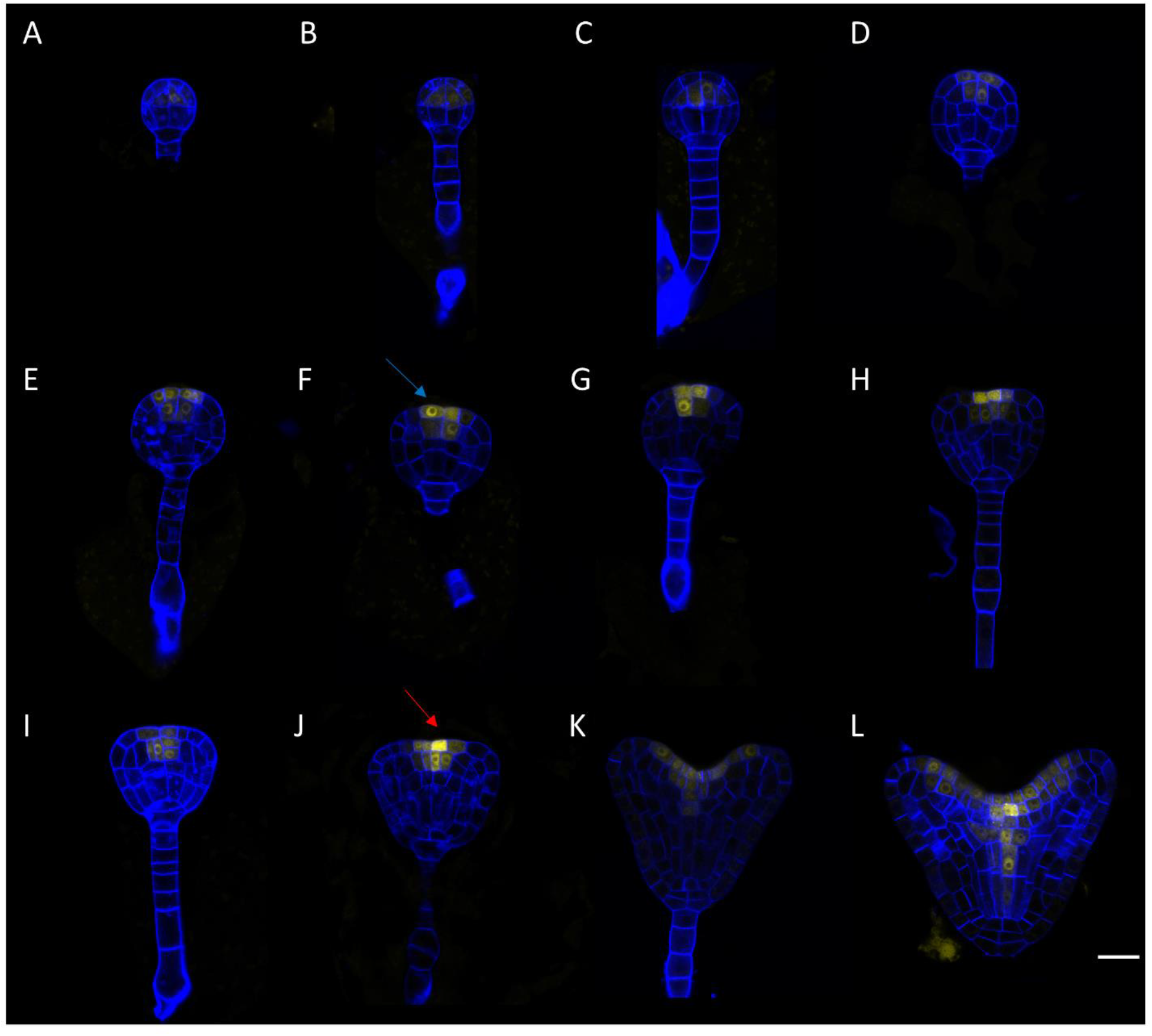
ZPR3 expression pattern in different stages of embryogenesis. Protein accumulation of ZPR3g:2xYPET-3xFLAG in the apical domain of the globular stage embryo, and in the adaxial side of the cotyledons, the vasculature and the meristem in the heart stage embryo. a-b, early globular. c-e, mid globular. f-g, late globular. h-i, transition. J, early heart. k-l, late heart. Red arrow points to cell with higher accumulation of ZPR3 protein in the cytosol (j). Blue arrow points to cell with higher accumulation of ZPR3 protein in the nucleus (f). Scale bar = 20 μm.

**Sup. Fig. 5.**
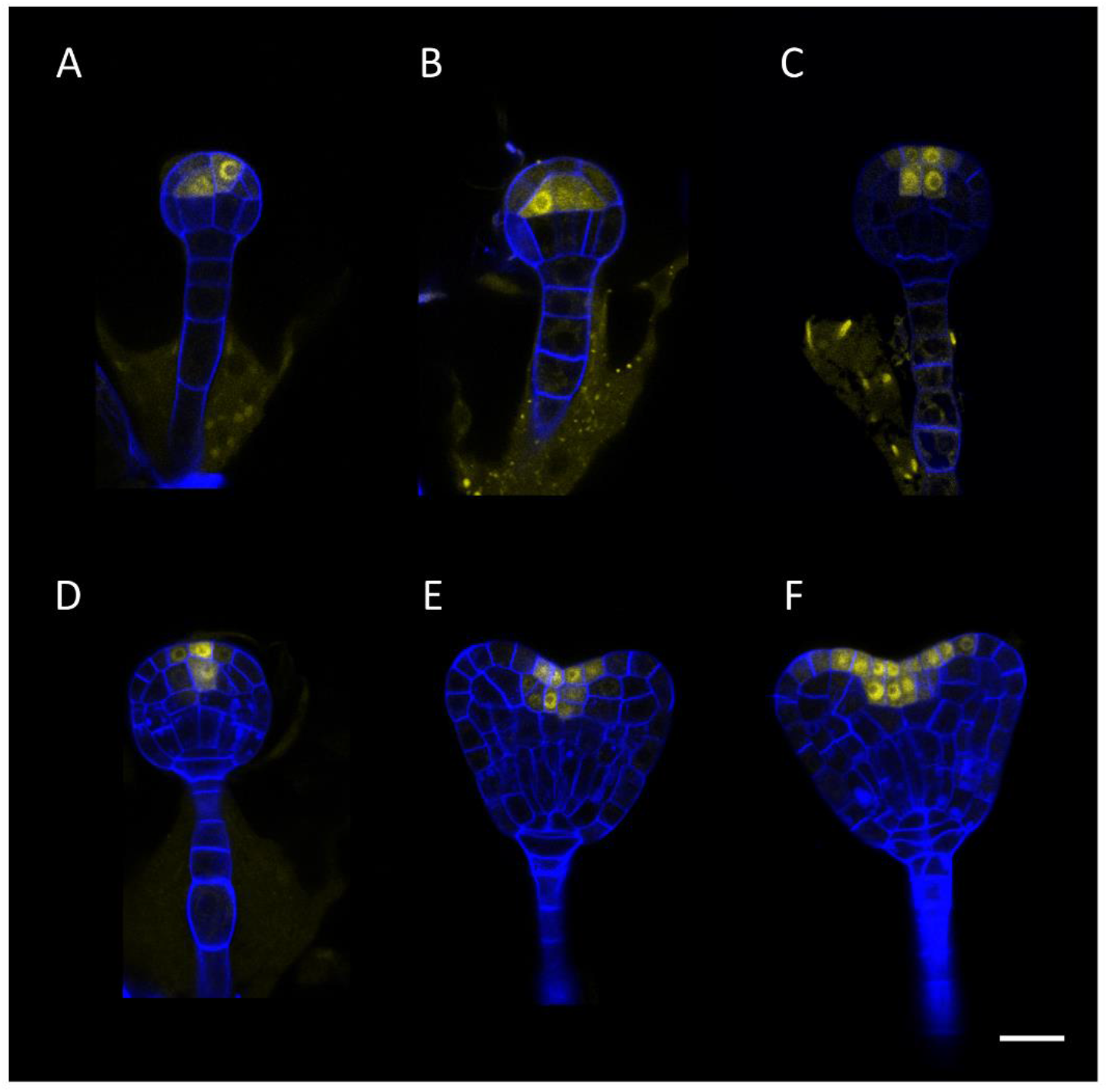
ZPR4 expression pattern in different stages of embryogenesis. Protein accumulation of ZPR4g:2xYPET-3xFLAG in the apical domain of the globular stage embryo, and in the adaxial side of the cotyledons, the vasculature and the meristem in the heart stage embryo. a, early globular. b, mid globular. c, late globular. d, transition. e-f, late heart. Scale bar = 20 μm.

**Sup. Fig. 6.**
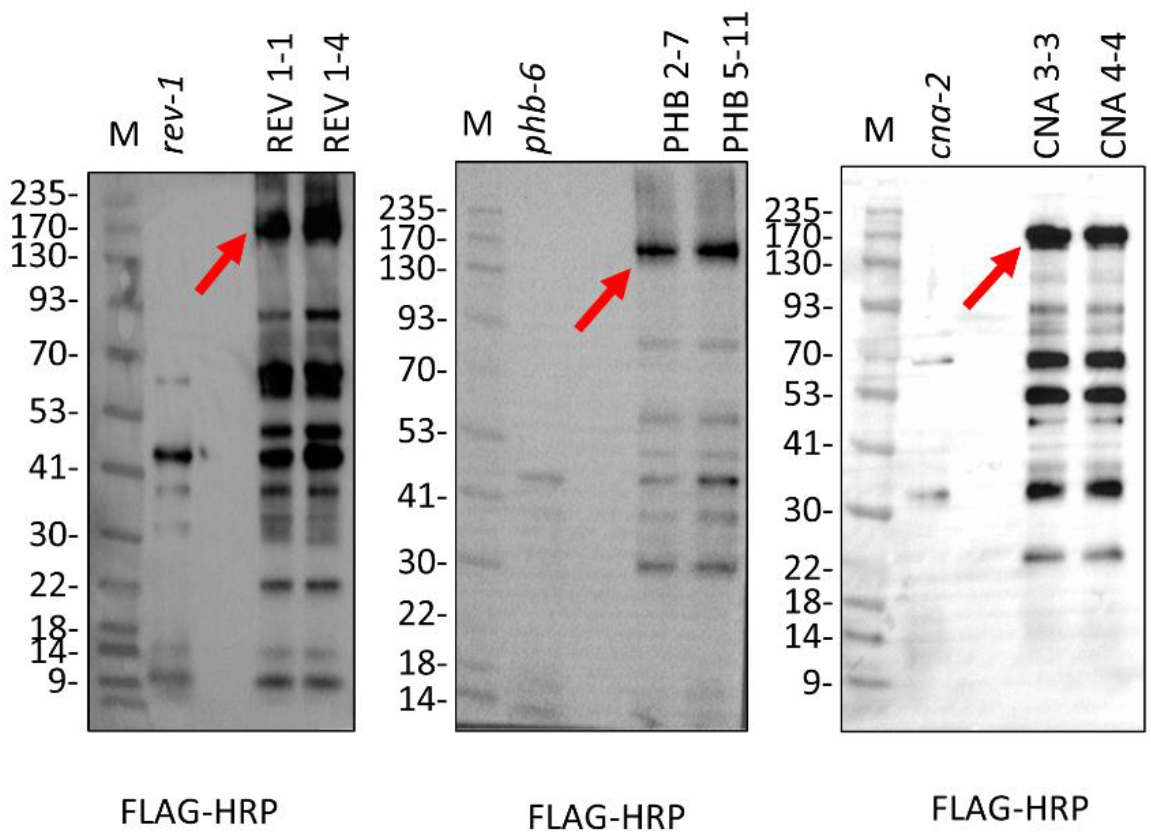
An anti-FLAG blots of the HD-ZIP III IPMS elution fraction. Red arrows point to an enrichment of the tagged HD-ZIP III protein in the two independent biological replicate lines in comparison to the *hd-zip III* mutant control.

## References

Alonso, J. M., & Stepanova, A. N. (2014). A recombineering-based gene tagging system for Arabidopsis. Bacterial Artificial Chromosomes: Second Edition, 233–243. https://doi.org/10.1007/978-1-4939-1652-8_11

Amoutzias, G. D., Robertson, D. L., Van de Peer, Y., & Oliver, S. G. (2008). Choose your partners: dimerization in eukaryotic transcription factors. In Trends in Biochemical Sciences (Vol. 33, Issue 5). https://doi.org/10.1016/j.tibs.2008.02.002

Bhatia, N., Bozorg, B., Larsson, A., Ohno, C., Jönsson, H., & Heisler, M. G. (2016). Auxin Acts through MONOPTEROS to Regulate Plant Cell Polarity and Pattern Phyllotaxis. Current Biology, 26(23). https://doi.org/10.1016/j.cub.2016.09.044

Brumos, J., Zhao, C., Gong, Y., Soriano, D., Patel, A. P., Perez-Amador, M. A., Stepanova, A. N., & Alonso, J. M. (2020). An Improved Recombineering Toolset for Plants. The Plant Cell, 32(1). https://doi.org/10.1105/tpc.19.00431

Byrne, M. E. (2006). Shoot meristem function and leaf polarity: The role of class III HD-ZIP genes. PLoS Genetics, 2(6), 0785–0790. https://doi.org/10.1371/journal.pgen.0020089

Caggiano, M. P., Yu, X., Bhatia, N., Larsson, A., Ram, H., Ohno, C. K., Sappl, P., Meyerowitz, E. M., Jönsson, H., & Heisler, M. G. (2017). Cell type boundaries organize plant development. ELife, 6. https://doi.org/10.7554/eLife.27421

Carlsbecker, A., Lee, J. Y., Roberts, C. J., Dettmer, J., Lehesranta, S., Zhou, J., Lindgren, O., Moreno-Risueno, M. A., Vatén, A., Thitamadee, S., Campilho, A., Sebastian, J., Bowman, J. L., Helariutta, Y., & Benfey, P. N. (2010). Cell signalling by microRNA165/6 directs gene dose-dependent root cell fate. Nature, 465(7296). https://doi.org/10.1038/nature08977

Chandler, J. W., Cole, M., Flier, A., Grewe, B., & Werr, W. (2007). The AP2 transcription factors DORNROSCHEN and DORNROSCHEN-LIKE redundantly control Arabidopsis embryo patterning via interaction with PHAVOLUTA. Development, 134(9), 1653–1662. https://doi.org/10.1242/dev.001016

Clough, S. J., & Bent, A. F. (1998). Floral dip: A simplified method for Agrobacterium-mediated transformation of Arabidopsis thaliana. Plant Journal, 16(6). https://doi.org/10.1046/j.1365-313X.1998.00343.x

Cox, J., & Mann, M. (2008). MaxQuant enables high peptide identification rates, individualized p.p.b.-range mass accuracies and proteome-wide protein quantification. Nature Biotechnology, 26(12). https://doi.org/10.1038/nbt.1511

Damodaran, S., Dubois, A., Xie, J., Ma, Q., Hindié, V., & Subramanian, S. (2019). GmZPR3d interacts with GmHD-ZIP III proteins and regulates soybean root and nodule vascular development. International Journal of Molecular Sciences, 20(4). https://doi.org/10.3390/ijms20040827

Deppmann, C. D., Acharya, A., Rishi, V., Wobbes, B., Smeekens, S., Taparowsky, E. J., & Vinson, C. (2004). Dimerization specificity of all 67 B-ZIP motifs in Arabidopsis thaliana: A comparison to Homo sapiens B-ZIP motifs. Nucleic Acids Research, 32(11). https://doi.org/10.1093/nar/gkh653

Donner, T. J., Sherr, I., & Scarpella, E. (2009). Regulation of preprocambial cell state acquisition by auxin signaling in Arabidopsis leaves. Development, 136(19). https://doi.org/10.1242/dev.037028

Duclercq, J., Assoumou Ndong, Y. P., Guerineau, F., Sangwan, R. S., & Catterou, M. (2011). Arabidopsis shoot organogenesis is enhanced by an amino acid change in the ATHB15 transcription factor. Plant Biology, 13(2). https://doi.org/10.1111/j.1438-8677.2010.00363.x

Elhiti, M., & Stasolla, C. (2009). Structure and function of homodomain-leucine zipper (HD-Zip) proteins. Plant Signaling & Behavior, 4(2), 86–88. https://doi.org/10.4161/psb.4.2.7692

Emery, J. F., Floyd, S. K., Alvarez, J., Eshed, Y., Hawker, N. P., Izhaki, A., Baum, S. F., & Bowman, J. L. (2003). Radial Patterning of Arabidopsis Shoots by Class III HD-ZIP and KANADI Genes. Current Biology, 13(20), 1768–1774. https://doi.org/10.1016/j.cub.2003.09.035

Eshed, Y., Baum, S. F., Perea, J. V., & Bowman, J. L. (2001). Establishment of polarity in lateral organs of plants. Current Biology, 11(16). https://doi.org/10.1016/S0960-9822(01)00392-X

Floyd, S. K., Ryan, J. G., Conway, S. J., Brenner, E., Burris, K. P., Burris, J. N., Chen, T., Edger, P. P., Graham, S. W., Leebens-Mack, J. H., Pires, J. C., Rothfels, C. J., Sigel, E. M., Stevenson, D. W., Neal Stewart, C., Wong, G. K. S., & Bowman, J. L. (2014). Origin of a novel regulatory module by duplication and degeneration of an ancient plant transcription factor. Molecular Phylogenetics and Evolution, 81. https://doi.org/10.1016/j.ympev.2014.06.017

Grigg, S. P., Galinha, C., Kornet, N., Canales, C., Scheres, B., & Tsiantis, M. (2009). Repression of Apical Homeobox Genes Is Required for Embryonic Root Development in Arabidopsis. Current Biology, 19(17). https://doi.org/10.1016/j.cub.2009.06.070

Husbands, A. Y., Aggarwal, V., Ha, T., & Timmermans, M. C. P. (2016). In planta single-molecule pull-down reveals tetrameric stoichiometry of HD-ZIPIII:LITTLE ZIPPER complexes. Plant Cell, 28(8). https://doi.org/10.1105/tpc.16.00289

Jami-Alahmadi, Y., Pandey, V., Mayank, A. K., & Wohlschlegel, J. A. (2021). A robust method for packing high resolution c18 rp-nano-hplc columns. Journal of Visualized Experiments, 2021(171). https://doi.org/10.3791/62380

Juarez, M. T., Kui, J. S., Thomas, J., Heller, B. A., & Timmermans, M. C. P. (2004). microRNA-mediated repression of rolled leaf1 specifies maize leaf polarity. Nature, 428(6978), 84–88. https://doi.org/10.1038/nature02363

Kidner, C. A., & Martienssen, R. A. (2004). Spatially restricted microRNA directs leaf polarity through ARGONAUTE1. Nature, 428(6978), 81–84. https://doi.org/10.1038/nature02366

Kim, J., Jung, J. H., Reyes, J. L., Kim, Y. S., Kim, S. Y., Chung, K. S., Kim, J. A., Lee, M., Lee, Y., Kim, V. N., Chua, N. H., & Park, C. M. (2005). microRNA-directed cleavage of ATHB15 mRNA regulates vascular development in Arabidopsis inflorescence stems. Plant Journal, 42(1), 84–94. https://doi.org/10.1111/j.1365-313X.2005.02354.x

Kim, Y. S., Kim, S. G., Lee, M., Lee, I., Park, H. Y., Pil, J. S., Jung, J. H., Kwon, E. J., Se, W. S., Paek, K. H., & Park, C. M. (2008). HD-ZIP III activity is modulated by competitive inhibitors via a feedback loop in Arabidopsis shoot apical meristem development. Plant Cell, 20(4). https://doi.org/10.1105/tpc.107.057448

Lindsey, B. E., Rivero, L., Calhoun, C. S., Grotewold, E., & Brkljacic, J. (2017). Standardized method for high-throughput sterilization of Arabidopsis seeds. Journal of Visualized Experiments, 2017(128). https://doi.org/10.3791/56587

Lucas, W. J., Groover, A., Lichtenberger, R., Furuta, K., Yadav, S. R., Helariutta, Y., He, X. Q., Fukuda, H., Kang, J., Brady, S. M., Patrick, J. W., Sperry, J., Yoshida, A., López-Millán, A. F., Grusak, M. A., & Kachroo, P. (2013). The Plant Vascular System: Evolution, Development and Functions. In Journal of Integrative Plant Biology (Vol. 55, Issue 4). https://doi.org/10.1111/jipb.12041

Magnani, E., & Barton, M. K. (2011). A Per-ARNT-sim-like sensor domain uniquely regulates the activity of the homeodomain leucine zipper transcription factor REVOLUTA in Arabidopsis. Plant Cell, 23(2). https://doi.org/10.1105/tpc.110.080754

Mallory, A. C., Reinhart, B. J., Jones-Rhoades, M. W., Tang, G., Zamore, P. D., Barton, M. K., & Bartel, D. P. (2004). MicroRNA control of PHABULOSA in leaf development: Importance of pairing to the microRNA 5??? region. EMBO Journal, 23(16), 3356–3364. https://doi.org/10.1038/sj.emboj.7600340

Mayer, K. F. X., Schoof, H., Haecker, A., Lenhard, M., Jürgens, G., & Laux, T. (1998). Role of WUSCHEL in regulating stem cell fate in the Arabidopsis shoot meristem. Cell, 95(6). https://doi.org/10.1016/S0092-8674(00)81703-1

McConnell, J. R., & Barton, M. K. (1998). Leaf polarity and meristem formation in Arabidopsis. Development, 125(15). https://doi.org/10.1242/dev.125.15.2935

McConnell, J. R., Emery, J., Eshed, Y., Bao, N., Bowman, J., & Barton, M. K. (2001). Role of PHABULOSA and PHAVOLUTA in determining radial patterning in shoots. Nature, 411(6838), 709–713. https://doi.org/10.1038/35079635

Moissiard, G., Bischof, S., Husmann, D., Pastor, W. A., Hale, C. J., Yen, L., Stroud, H., Papikian, A., Vashisht, A. A., Wohlschlegel, J. A., & Jacobsen, S. E. (2014). Transcriptional gene silencing by Arabidopsis microrchidia homologues involves the formation of heteromers. Proceedings of the National Academy of Sciences of the United States of America, 111(20). https://doi.org/10.1073/pnas.1406611111

Mukherjee, K., & Bürglin, T. R. (2006). MEKHLA, a novel domain with similarity to PAS domains, is fused to plant homeodomain-leucine zipper III proteins. Plant Physiology, 140(4). https://doi.org/10.1104/pp.105.073833

Nagata, K., & Abe, M. (2021). The lipid-binding START domain regulates the dimerization of ATML1 via modulating the ZIP motif activity in Arabidopsis thaliana. Development Growth and Differentiation, 63(8). https://doi.org/10.1111/dgd.12753

Otsuga, D., DeGuzman, B., Prigge, M. J., Drews, G. N., & Clark, S. E. (2001). REVOLUTA regulates meristem initiation at lateral positions. Plant Journal, 25(2), 223–236. https://doi.org/10.1046/j.1365-313X.2001.00959.x

Prigge, M. J., Otsuga, D., Alonso, J. M., Ecker, J. R., Drews, G. N., & Clark, S. E. (2005). Class III homeodomain-leucine zipper gene family members have overlapping, antagonistic, and distinct roles in Arabidopsis development. The Plant Cell, 17(1), 61–76. https://doi.org/10.1105/tpc.104.026161.1

Ramachandran, P., Carlsbecker, A., Etchells, J. P., & Turner, S. (2016). Class III HD-ZIPs govern vascular cell fate: An HD view on patterning and differentiation. Journal of Experimental Botany, 68(1), 55–69. https://doi.org/10.1093/jxb/erw370

Riechmann, J. L., Heard, J., Martin, G., Reuber, L., Jiang, C. Z., Keddie, J., Adam, L., Pineda, O., Ratcliffe, O. J., Samaha, R. R., Creelman, R., Pilgrim, M., Broun, P., Zhang, J. Z., Ghandehari, D., Sherman, B. K., & Yu, G. L. (2000). Arabidopsis transcription factors: Genome-wide comparative analysis among eukaryotes. Science, 290(5499). https://doi.org/10.1126/science.290.5499.2105

Schindele, P., & Puchta, H. (2020). Engineering CRISPR/LbCas12a for highly efficient, temperature-tolerant plant gene editing. Plant Biotechnology Journal, 18(5). https://doi.org/10.1111/pbi.13275

Sessa, G., Steindler, C., Morelli, G., & Ruberti, I. (1998). The Arabidopsis Athb-8, −9 and −14 genes are members of a small gene family coding for highly related HD-ZIP proteins. Plant Molecular Biology, 38(4). https://doi.org/10.1023/A:1006016319613

Smith, Z. R., & Long, J. A. (2010). Control of Arabidopsis apical–basal embryo polarity by antagonistic transcription factors. Nature, 464(7287), 423–426. https://doi.org/10.1038/nature08843

Somssich, M., Je, B. Il, Simon, R., & Jackson, D. (2016). CLAVATA-WUSCHEL signaling in the shoot meristem. In Development (Cambridge) (Vol. 143, Issue 18). https://doi.org/10.1242/dev.133645

Soyars, C. L., James, S. R., & Nimchuk, Z. L. (2016). Ready, aim, shoot: Stem cell regulation of the shoot apical meristem. In Current Opinion in Plant Biology (Vol. 29). https://doi.org/10.1016/j.pbi.2015.12.002

Talbert, P. B., Adler, H. T., Parks, D. W., & Comai, L. (1995). The REVOLUTA gene is necessary for apical meristem development and for limiting cell divisions in the leaves and stems of Arabidopsis thaliana. Development (Cambridge, England), 121(9), 2723–2735. http://www.ncbi.nlm.nih.gov/pubmed/7555701

Thomas, P. D., Campbell, M. J., Kejariwal, A., Mi, H., Karlak, B., Daverman, R., Diemer, K., Muruganujan, A., & Narechania, A. (2003). PANTHER: A library of protein families and subfamilies indexed by function. Genome Research, 13(9). https://doi.org/10.1101/gr.772403

Warming, S., Costantino, N., Court, D. L., Jenkins, N. A., & Copeland, N. G. (2005). Simple and highly efficient BAC recombineering using galK selection. Nucleic Acids Research, 33(4). https://doi.org/10.1093/nar/gni035

Weits, D. A., Kunkowska, A. B., Kamps, N. C. W., Portz, K. M. S., Packbier, N. K., Nemec Venza, Z., Gaillochet, C., Lohmann, J. U., Pedersen, O., van Dongen, J. T., & Licausi, F. (2019). An apical hypoxic niche sets the pace of shoot meristem activity. Nature, 569(7758). https://doi.org/10.1038/s41586-019-1203-6

Wenkel, S., Emery, J., Hou, B.-H., Evans, M. M. S., & Barton, M. K. (2007). A Feedback Regulatory Module Formed by LITTLE ZIPPER and HD-ZIPIII Genes. The Plant Cell Online, 19(11), 3379–3390. https://doi.org/10.1105/tpc.107.055772

Williams, L., Grigg, S. P., Xie, M., Christensen, S., & Fletcher, J. C. (2005). Regulation of Arabidopsis shoot apical meristem and lateral organ formation by microRNA miR166g and its AtHD-ZIP target genes. Development, 132(16). https://doi.org/10.1242/dev.01942

Xie, Z., Allen, E., Fahlgren, N., Calamar, A., Givan, S. A., & Carrington, J. C. (2005). Expression of Arabidopsis MIRNA genes. Plant Physiology, 138(4). https://doi.org/10.1104/pp.105.062943

Xu, Q., Li, R., Weng, L., Sun, Y., Li, M., & Xiao, H. (2019). Domain-specific expression of meristematic genes is defined by the LITTLE ZIPPER protein DTM in tomato. Communications Biology, 2(1). https://doi.org/10.1038/s42003-019-0368-8

Yadav, R. K., Perales, M., Gruel, J., Girke, T., Jönsson, H., & Venugopala Reddy, G. (2011). WUSCHEL protein movement mediates stem cell homeostasis in the Arabidopsis shoot apex. Genes and Development, 25(19). https://doi.org/10.1101/gad.17258511

Yan, L., Wei, S., Wu, Y., Hu, R., Li, H., Yang, W., & Xie, Q. (2015). High-Efficiency Genome Editing in Arabidopsis Using YAO Promoter-Driven CRISPR/Cas9 System. In Molecular Plant (Vol. 8, Issue 12). https://doi.org/10.1016/j.molp.2015.10.004

Zhou, G. K., Kubo, M., Zhong, R., Demura, T., & Ye, Z. H. (2007). Overexpression of miR165 affects apical meristem formation, organ polarity establishment and vascular development in Arabidopsis. Plant and Cell Physiology, 48(3), 391–404. https://doi.org/10.1093/pcp/pcm008

Zhu, H., Hu, F., Wang, R., Zhou, X., Sze, S. H., Liou, L. W., Barefoot, A., Dickman, M., & Zhang, X. (2011). Arabidopsis argonaute10 specifically sequesters miR166/165 to regulate shoot apical meristem development. Cell, 145(2), 242–256. https://doi.org/10.1016/j.cell.2011.03.024

